# Contact-number-driven virus evolution: a multi-level modeling framework for the evolution of acute or persistent RNA virus infection

**DOI:** 10.1101/2022.12.22.521662

**Authors:** Junya Sunagawa, Ryo Komorizono, William S. Hart, Robin N. Thompson, Akiko Makino, Keizo Tomonaga, Shingo Iwami, Ryo Yamaguchi

## Abstract

Viruses evolve in infected host populations, and host population dynamics affect viral evolution. RNA viruses with a short duration of infection and a high peak viral load, such as and SARS-CoV-2, are maintained in human populations. By contrast, RNA viruses characterized by a long infection duration and a low peak viral load (e.g., borna disease virus) can be maintained in nonhuman populations, and why the persistent viruses evolved has been rarely explored. Here, using a multi-level modeling approach including both individual-level virus infection dynamics and population-scale transmission, we consider virus evolution based on the host environment, specifically, the effect of the contact history of infected hosts. We found that, with a highly dense contact history, viruses with a high virus production rate but low accuracy are likely to be optimal, resulting in a short infectious period with a high peak viral load. In contrast, with a low-density contact history, viral evolution is toward low virus production but high accuracy, resulting in long infection durations with low peak viral load. Our study sheds light on the origin of persistent viruses and why acute viral infections but not persistent virus infection tends to prevail in human society.

## Text

Because they lack proofreading activity in their RNA-dependent RNA polymerase, RNA viruses exhibit extremely high mutation rates, generating 10^-6^ to 10^-4^ substitutions per nucleotide, orders of magnitude greater than those of most DNA-based life forms [1–3]. The accuracy (or inaccuracy) of the polymerase is the source of intra-host viral diversity and determines viral propagation and virulence *in vivo* [4, 5].

In general, acute infectious RNA viruses, such as SARS-CoV-2, are highly proliferative and release large numbers of progeny viral particles [6]. From an epidemiologic point of view, acute infectious RNA viruses are highly pathogenic [7, 8]. Infections with most non-retroviral RNA viruses result in the characteristic symptoms and signs of acute illness. On the other hand, persistent infection is a different strategy. The properties of RNA viruses vary widely, among which non-retroviral RNA viruses can often be broadly classified into acute infection and persistent (chronic) infection. Some non-retroviral RNA viruses, such as hepatitis C virus, coxsackievirus, and borna disease virus, establish persistent infection and replicate in the host for long periods [9]. Whereas acute infectious viruses are eliminated by the immune response within a few weeks, persistent infectious viruses are maintained in the host for at least several months or even years. In most cases, patients are mildly symptomatic or asymptomatic during persistent infection. Because it is not easy to discern the causes of mortality in the course of a viral infection, given the multiple viral and host factors involved in viral pathogenesis, but many acutely infectious RNA viruses cause fatal disease soon after infection (e.g. Rabie virus, Nipah virus), while some persistent infectious RNA viruses cause nonfatal disease after infection (e.g. Borna disease virus, Pegi virus) [10]. The persistent infectious virus can coexist with the host by reducing the amount of progeny virus released from the infected cells, slowing down the replication rate, decreasing evolutionary rate, and evading the host’s immune system to use the host as its own reservoir until the next opportunity to infect [11–15]. These different viral phenotypes of acute and persistent infection express different strategies in the host population, ranging from intracellular replication rates and innate immune response to interindividual transmission dynamics [9].

For gaining a better understanding of how viral properties is optimized, the lifestyles of the host organisms are of critical importance. Different host behaviors provide different environments, with some environments favoring particular viral characteristics. Human activities in the Anthropocene have greatly affected the global environment, increasing the frequency of human-animal contact and various human-induced evolutionary changes in animals [16]. Traditionally, it is considered that acute infection evolved because of population-level transmission potential, that is, the potential for many susceptible hosts to come into contact with an infected individual [17]. This rose a question of “why did persistent infectious RNA viruses evolve?”. Commonly, the population-level transmission potential is characterized by the reproduction number [8]. However, the environmental conditions in the aspect of contact history that favor the evolution of persistent infectious RNA viruses have not yet been explored. In addition, in terms of the within-host virus life cycle, it is unclear how viral properties such as the proliferation rate and the accuracy of viral replication affect virus evolution under different environmental conditions. Another unresolved issue is why acute infections are generally more prevalent than persistent infections in the current profile of viral diversity in human populations [11].

To tackle these challenges, we employed a multi-level modeling approach including both individual-level virus infection dynamics and population-scale transmission. To seek optimal strategy across a broad range of viruses rather to explain properties of a specific virus, we simply assume that an optimization occurs at an epidemiological equilibrium (i.e., focusing on equilibrium solutions). Although elaborate discussions are possible by considering *R*_0_ and balance with other high level population criterion like lifespan of the host in models specific to each virus (e.g., [18, 19]), we examine the long-term equilibrium evolutionary outcome in the context of stable endemic diseases which solely maximizes basic reproductive number, *R*_0_. Our simplification is valid unless considering the situation where pathogens often showing non-equilibrium dynamics such as antigenic escape from host immune defenses through evolution [20]. Under different environments characterized by different contact numbers in populations, we evaluated how measures of the proliferation rate and accuracy in the viral life cycle affect population-level transmission potential and therefore RNA virus evolution.

## Results

### Virus evolution in different contact scenarios

To explore how virus evolution is influenced by contact number, we first developed a probabilistic multi-level model to characterize population-level transmission potential based on contact histories. Using a genetic algorithm (see **Materials and Methods** for further details), we investigated the evolution of the proliferation rate and the accuracy of replication in the viral life cycle, mainly reflecting into *p* and *ε*, respectively. We note *ε* includes multiple-factors such as the accuracy of virion assembly, the rate of virus particle aggregation and others, in addition to the accuracy of replication which is the main factor (see [Eqs.(1–4)]). From the assumption that the virus is optimized to increase its transmission potential, denoted by *R*_0_, the genetic algorithm heuristically searches for the optimal *R*_0_ as a population-based method with a group of solutions made up of combinations of *p* and *ε*. Our model [Eqs.(7–8)] calculates *R*_0_ as the sum of secondary cases generated throughout the infectious period by a primary case (see **Fig. 1A**, **Materials and Methods** in detail). This means that the long-term equilibrium evolutionary outcome is investigated here. The durations of the infectious periods *D*(*V*_total_) in each parameter combination (i.e., *p* and *ε*) are calculated by Eqs.(5–6) based on the virus infection model [Eqs.(1–4)] with other fixed parameters (see **Table S1**). The calculated infectious period for each parameter pair is described in **Fig.1B**.

**Fig. 1.**
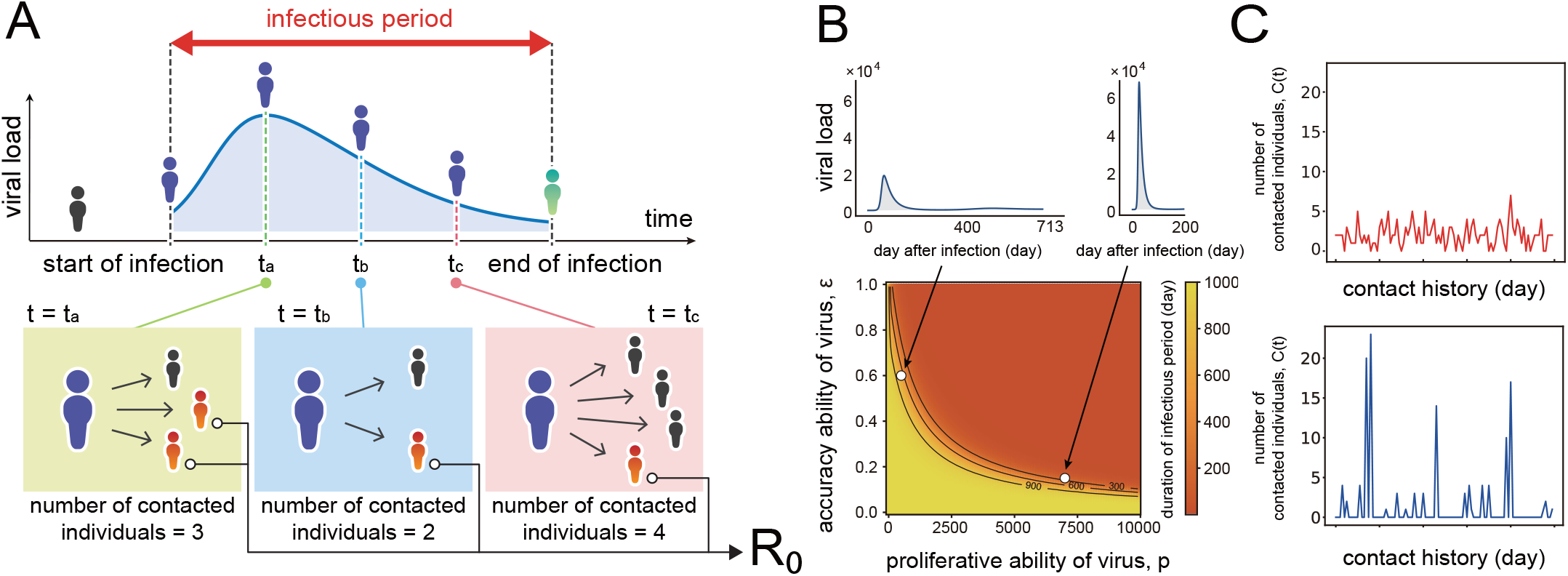
Schematic illustration of multi-level disease transmission. A schematic of our model is depicted in **(A)**. Virus transmission occurs from an infected individual to susceptible individuals depending on the infected individual’s infectivity, which depends on their viral load. At each time step (i.e., day), the focal infected individual has contacts with multiple susceptible individuals. Here the contact numbers are assumed to follow a negative binomial distribution, which do not depend on the viral load. The sum of newly infected individuals (i.e., [number of contacted individuals per day] × [probability of infection per contacted individual]) during the infectious period is calculated as *R*_0_. The duration of the infectious periods for each parameter combination (*p* and *ε*) characterizing the proliferative ability and accuracy of viruses are calculated in **(B)** based on the virus infection model. The left panel corresponds to an infectious period of 713 days with a relatively low viral load, and the right panel represents an infectious period of 200 days with a high viral load. Examples of generated daily contact numbers characterizing environmental conditions are shown in **(C)**. The contact numbers under a negative binomial distribution with *k* = 100 and *θ* = 0.03 (the mean and variance are 3.0 and 3.1) and *k* = 0.12 and *θ* = 10 (the mean and variance are 1.2 and 13.3) are shown in the top and bottom panels, respectively.

To evaluate how the virus is optimized for different contact numbers (i.e., environments), we made two categories of contact history with daily numbers of contacts sampled from the negative binomial distribution (see **Materials and Methods** in detail). The first category is one with large mean and small variance, in which the contact number is always around the mean throughout contact the history and does not decrease too low or increase too high (**Fig.1C**, top). The other has small mean and large variance, in which the contact number is usually 0 but is sometimes very high (**Fig.1C**, bottom). For contact numbers with large mean and small variance, *k* = 100 and *θ = 0.03* (the mean number is 3) are fixed to generate the contact history. In the history with small mean and large variance, *k* = 0.12 and 0 = 10 (the mean number is 1.2). Here *k* and *θ* represent the shape parameter and scale parameter of the negative binomial distribution, respectively.

Given a contact history generated from the negative binomial distribution, as described above, we searched for the optimal point in the fitness landscape (i.e., the largest *R*_0_), that is, the optimal combination of *p* and *ε*, by using a genetic algorithm for each contact number (see **Materials and Methods** in detail). For the contact history with large mean and small variance (i.e., high-density contact history: (*k,θ*) = (100,0.03), **Fig.1C**, top), optimal fitness involves high virus production but low accuracy, which results in a short duration of infectious period with high peak viral load (we call this the *acute infection* phenotype) (**Fig.2A** and **Fig.S1**). In contrast, for the contact history with small mean and large variance (i.e., low-density contact history: (*k,θ) =* (0.12,10), **Fig.1C**, bottom), the virus is more likely to be optimized toward low virus production but high accuracy, resulting in a long infection duration with low peak viral load (we call this the *persistent infection* phenotype) (**Fig.2C** and **Fig.S1**). The white dots in **Fig.2A** and **Fig.2C** correspond to the coordinates of the combination of *p* and *ε* that maximize *R*_0_ with a given contact pattern.

**Fig. 2.**
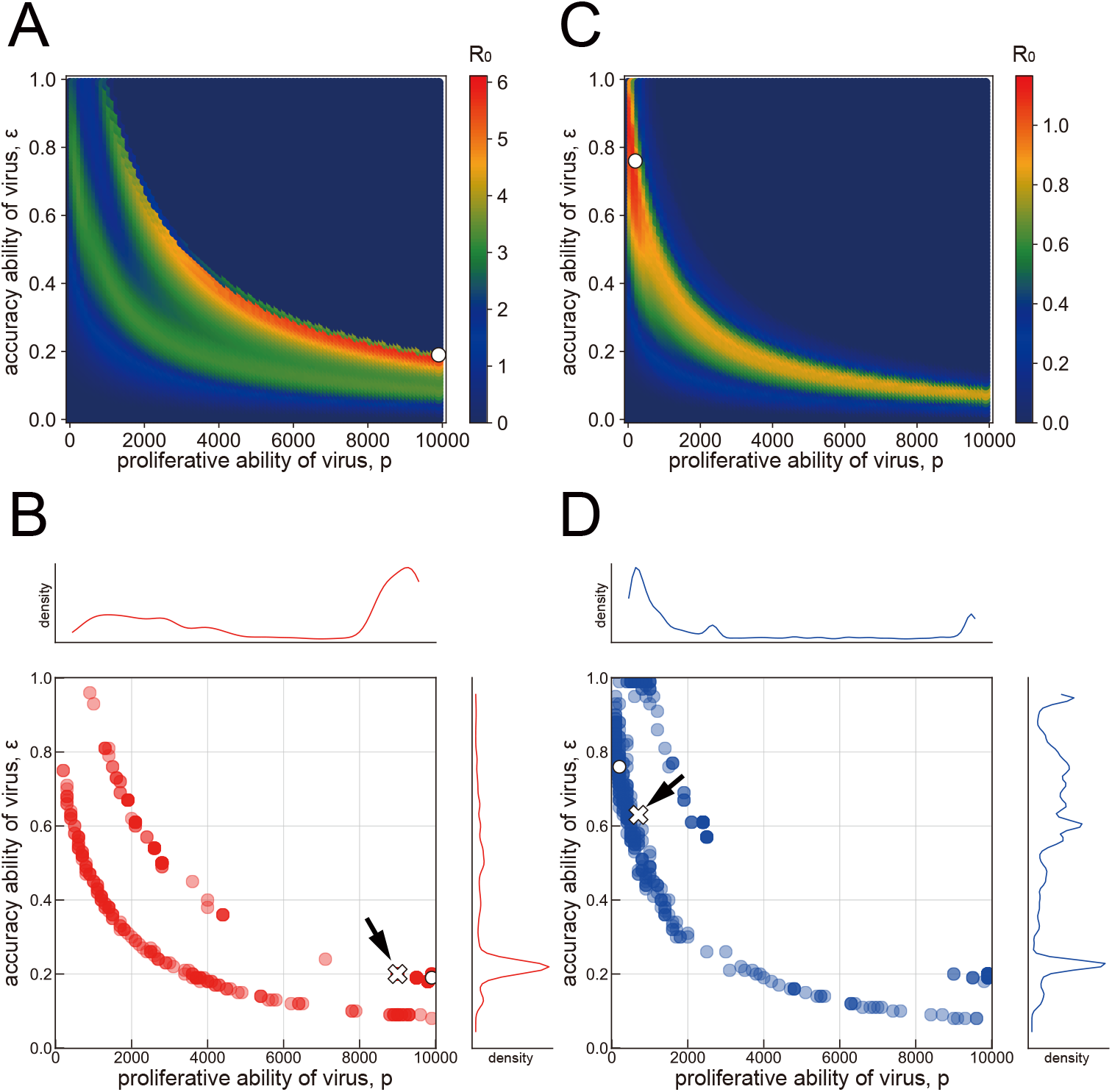
Virus evolution in a given environment. The optimal parameter combinations (*p* and *ε*) that maximize *R*_0_ are calculated for a parameter set of *k = 100* and *θ = 0.03* in **(A)**. The white dot represents the optimal point of *R*_0_ in a single simulation. Using the same parameter set, 500 optima based on independent contact histories are calculated in **(B)**. The kernel plots illustrated alongside the top and right parts are the marginal distributions of *p* and *ε*, respectively. The white dot exactly corresponds to the optimal point shown in a single simulation in **(A)**. The white cross indicated by the black arrow corresponds to the median of optimized parameter set of *p* and *ε* based on the 500 simulation runs. In a similar manner, the parameter set of *k* = 0.12 and *θ* = 10 is used to calculate the optimal parameter combinations (*p* and *ε*) in **(C)**. Using the same parameter sets, 500 evolutionary consequences for independently drawn contact histories are shown in **(D)**. The white dot and white cross correspond to the optimal point of the parameter set and the median point, respectively.

With 500 independent iterations of this process with the same parameter sets, the evolutionary endpoint of each iteration is distributed as shown in **Fig.2B** and **Fig.2D**. The white dots in **Fig.2B** and **Fig.2D** correspond to the evolutionary endpoint in each single simulation run, **Fig.2A** and **Fig.2C**, respectively. The white crosses indicated by the black arrows in **Fig.2B** and **Fig.2D** correspond to the median parameter set of *p* and *ε* after evolution under the given environment sequence of the contact history.

In addition to these contact histories, we considered the evolution of viral phenotype under other contact histories in scale-free networks and small-world networks. In the analysis on networks, we interpret the evolutionary consequences of the phenotypes similarly. Under networks where hubs exist or where individuals are in contact with each other evenly and with high frequency, the average value of the contact history is stable at a high value, and the acute infection type is likely to be optimal, and vice versa (**Fig. S2** and **S3**). As described in the next section, we investigated how the trend of virus evolution changes under the different environments.

### Virus evolution depending on the difference in mean contact number

We explored the optimal properties of the virus (proliferation rate and accuracy of replication, *p* and *ε*, respectively) for transmission by gradually changing the mean number of daily contacts in the contact history. To represent various contact histories, we explored a total of 441 patterns of contact histories ranging from 0.01 to 100. That is, we used the following parameter sets: shape parameter *k* ranges from 10^-1.0^, 10^-0.9^, 10^-0.8^,..., 10^0.9^, 10^1.0^ (sum up to 21 points), as is also the case with scale parameter *θ*, and each contact history is made by the combination of each *k* and *θ*. We conducted a total of 500 independent iterations for each parameter set of (*k, θ*) and determined the evolutionary consequences of the median parameter set of *p* and *ε*.

We found that viruses are optimized toward low production but high accuracy (the persistent infection phenotype) in regions of low-density contact history (the mean contact number from 0.01 to 1) regardless of the variance in daily contacts (white dots 3 and 4 in **Fig.3A** and **Fig.3B**, and **Fig.S4**). As discussed above, these optimized virus infection dynamics showed a longer duration of infectious period with low peak viral load (white dots 3 and 4 in **Fig.3C** and **Fig.3D**). In contrast, in the region of a highly dense contact history ranging from 30 to 100, viruses were optimized with high production but low accuracy (white dots 1 and 2 in **Fig.3A** and **Fig.3B**), representing the acute infection phenotype with the shorter duration of infectious period and high peak viral load (white dots 1 and 2 in **Fig.3C** and **Fig.3D**). With respect to the transmission potential, *R*_0_, a logarithmic increase is observed as the mean contact number increases (**Fig.3E**). Note that some of the persistent infection phenotypes evolving under the low-density contact history (white dot 4 in **Fig.3A, 3B, 3D**, and **3E**) might have a low transmission potential characterized by *R*_0_ <1 (see **Discussion**).

**Fig. 3.**
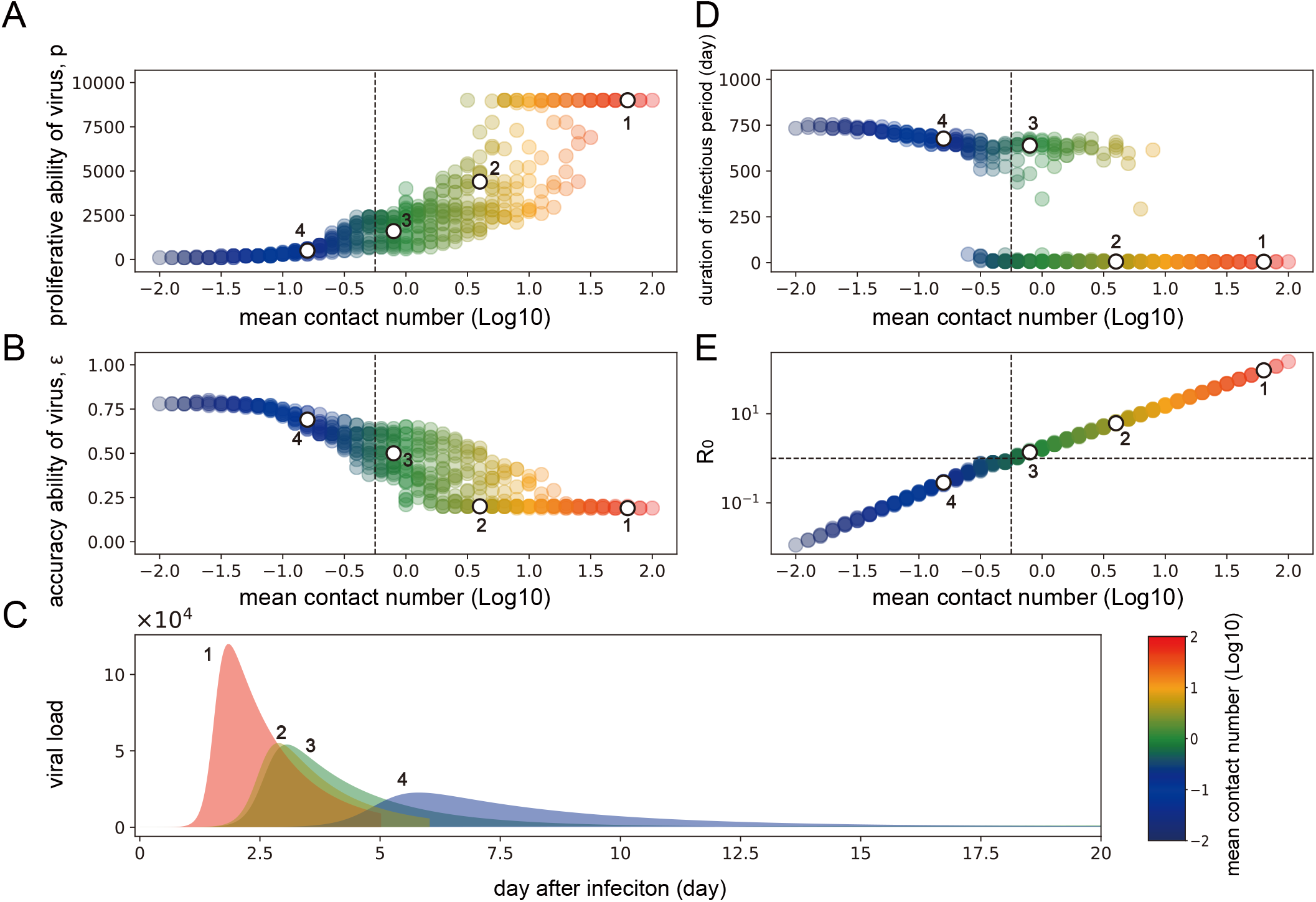
Virus evolution in various scenarios of contact history. A total of 441 contact history patterns are explored to determine the viral evolution to an optimal *R*_0_ in a given environmental scenario. 500 iterations of each parameter combination of *k* and *θ* are conducted to determine the optimized parameter set of *p* and *ε* characterizing final virus infection dynamics (i.e., different viral infection phenotypes). The median of the optimized parameter set of *p* and *ε* based on the 500 simulation runs is shown against the mean contact number of histories (with logarithmic increase) determined by each parameter set of *k* and *θ*. The combination of *p* and *ε* is plotted in **(A)** and **(B)**, separately. Colors represent the degree of the mean contact number: blue corresponds to a low mean contact number, whereas red corresponds to a high mean contact number. How virus infection dynamics vary with the mean contact number is shown in (**C**). Several examples of the virus infection dynamics are shown to represent the difference in the infectious period and the peak viral load. Numbers 1, 2, 3, and 4 in **(C)** are the corresponding dynamics illustrated as white dots in **(A)**, **(B)**, **(D)**, and **(E)**, respectively. The duration of the infectious period and *R*_0_ varied with the mean contact number are plotted in (**D**) and (**E**), respectively. In **(E)**, the vertical dashed line separates the area of mean contact number by an index, *R*_0_ = 10^0^(= 1). The corresponding dashed line is also illustrated in **(A)**, **(B)**, and **(D)**.

Importantly, we demonstrated that, if the contact history is of intermediate density (with mean contact numbers ranging from 1 to 30), viruses can be optimized toward either acute or persistent infection, showing a critical transition on the virus infection phenotypes (**Fig.3D**). This bistable-like pattern may appear by chance owing to the stochasticity of contact number history, even when drawn from the same distribution. Further discussion is provided below.

## Discussion

In this study, we investigated the impact of the proliferation rate and the accuracy of replication on viral evolution by considering optimal combinations for transmission under different contact histories of a host organism. Specifically, assuming an epidemiological equilibrium, we developed a probabilistic multi-level model that characterizes population-level transmission potential based on individual-level virus infection dynamics. As with many evolutionary biological models, our general model can be applied across many taxa to interpret effects on the diversity of shape of viral load. It should be noted that some studies have already existed to address within- and between host(s) dynamics. There are, however, several differences between our model and previous ones. First, to our knowledge, none of previous researches assumed the relationship between viral load and viral duration of infection (e.g., [19, 21]). We explicitly defined the duration of infectious period as a function of viral load (Eqs.(5–6)). Second, researches considering contact histories as an arena of virus evolution didn’t assume stochasticity [22] or extreme rare events [21]. We adopted a negative binomial distribution to generate various contact histories, which corresponds to a broad range of host species’ behavior. Using the current multi-level model, we especially focused on the condition where persistent viruses evolve.

The acute and persistent infection phenotypes are classified in terms of the duration of the infectious period and the peak viral load, which is determined by the combination of the accuracy of genome RNA replication, *ε*, and its production rate, *p*. The acute infection phenotype (with a short infectious period and high peak viral load) is achieved in the parameter range where the production rate of virus particles is high and the replication accuracy is low. Although our model does not explicitly include the mutation process by which new viral phenotypes are generated (cf. [21]), this corresponds to a scenario with a low accuracy of replication in the current model, which is the case for influenza viruses with around 90% of noninfectious activities [23, 24]. On the contrary, for the persistent infection phenotype, a long infectious period with a low peak viral load is obtained when there is high precision of virus replication and a low rate of replication. The properties characterized in this parameter region correspond to pathogens such as the borna virus [25, 26].

Our main finding is that the acute infection phenotype is optimal under conditions of a contact history involving frequent contacts between hosts (**Fig. 2AB** and **Fig. 3**). In this environment, the virus does not need to stay in the infected host for a long time, because the focal infected individual can quickly obtain another opportunity to infect (called a “leaving-home” strategy in [27]). Therefore, an effective strategy is to sustain a high peak viral load and adopt an increased probability of infection even for a short period of time (**Fig. S1**). On the other hand, in an environment with a small mean and a large variance in contact history, since there is a possibility that no contact will occur at all in a short period of infection (**Fig. 2CD** and **Fig. 3**), the virus must stay in the infected host for a long time (called the “stay-at-home” strategy in [27]). Hence, *R*_0_ increases by maintaining an infectious period with a long duration while suppressing the peak viral load (**Fig. S1**). This corresponds to the evolution of the persistent infection phenotype. Thus, the contact history of infected hosts determines the direction of viral evolution balanced between the duration of infectious period and peak viral load. That is, persistent-acute infection phenotype switching is induced depending on the environment.

In this study, one of the parameters that influenced the infection phenotype was the replication accuracy *ε*. The RNA-dependent RNA polymerase encoded by RNA viruses lacks proofreading activity, unlike DNA viruses. Owing to this low fidelity of viral polymerase, the genome sequence exists as diversified “quasispecies” in infected individuals [28–30]. These intra-host quasispecies are involved in viral fitness *in vivo* and affect the virulence and transmission ability of a virus [4, 28, 31]. Interestingly, low-fidelity mutants of coxsackievirus B3, which primarily infects the heart muscle, have lost the ability to establish persistent infection *in vivo* [14]. This result on the relationship between the fidelity of viral polymerase and persistent infection *in vivo* are consistent with our study and support the model that replication accuracy *ε* determines the establishment of the persistent infection phenotype (see supplementary material, Text S1 for detailed discussions). Interestingly, pandemic noroviruses of the GII.4 lineage have an approximately 1.7-fold higher mutation rate than the other strains [32]. In DNA viruses, there are many known persistently infectious viruses such as herpes simplex virus type 2, Epstein-Barr virus and cytomegalovirus while most non-retroviral RNA viruses are acutely infectious [11]. This may be due to the fast evolutionary rate unique to RNA viruses and host population density, which are drivers of viral evolution and may cause phenotypic differences between them.

From an evolutionary perspective, viruses adapt to the environments they experienced in the past, and their traits are optimized accordingly. Our exhaustive simulations with the multi-level model suggest that the acute infection phenotype is more likely to be optimal when individuals live in high densities or when there is a highly connected network of hosts (as is often the case for humans and other social animals). On the other hand, the persistent infection phenotype may be optimal in organisms that do not form groups and where the environmental variance for the virus is largely dependent on their life history; for example, if the contact rate increases only during the breeding season. The unpredictability of the presence of any individual, which is referred to “biotic drift,” can influence evolutionary consequences [33].

Our simulations also show that the evolutionary endpoints of *R*_0_ are larger in the acute infection phenotype than in the persistent infection phenotype (**Fig. 3E**). For example, with low-density contact histories ranging 0.01 to 0.78, *R*_0_ decreases below 1, meaning that viruses with the persistent infection phenotype are unlikely to be sustained from the perspective of population-level transmission, even though the virus phenotype adapts to the low-density contact histories. This implies that acute virus infection, such as with SARS-CoV-2, will easily prevail in human society. Borna disease virus, an RNA virus that can establish persistent infection without cytopathic effects for a long time, has been observed as an endemic disease in Germany but has not reached a large-scale epidemic [34–36]. On the other hand, even if viruses with the persistent infection phenotype are not transmitted among individuals in the context of low-density contact history, an increase in contact numbers may lead an optimized virus with the acute infection phenotype, increasing a high transmission potential within a host population. In fact, many acute infectious viruses that infect humans have been proposed to have been optimized through continuous mass infection after the formation of highly dense human communities for cultural reasons, such as for agriculture [37–40].

Large-scale genome sequencing and phylogenetic analysis of SARS-CoV-2 in the epidemic furthers the conclusions of this study. Epidemiological studies have shown that SARS-CoV-2 evolved from pre-alpha to alpha, beta and delta variants via host-to-host transmission [41]. These variants of concern (VOCs) transmit more quickly in host population compared to background, suggesting that they may have shorter intrinsic generation times [42]. Moreover, the molecular clock modeling using the GISAID database and Bayesian analysis suggested that the substitution rate of SARS-CoV-2 is increasing with the emergence of VOCs relative to the background [43]. In the above, it can be inferred that the continuous chain of infection within the host population facilitates viral evolution and contributes to the establishment of further acutely infectious viral phenotypes.

Going forward, substantial changes in contact patterns between hosts could lead to changes in the characteristics of emerging viruses with pandemic potential. Increasing population sizes and densities, and higher rates of global travel, lead to larger numbers of contacts between individuals. Changes in land use are leading to more frequent contacts between humans and potential zoonotic reservoirs. This not only increases the risk for emergence of pathogens with pandemic potential [44], but also, as we have shown, affects the characteristics of the viruses that successfully emerge. Specifically, higher densities of contacts drive evolution of viruses with higher viral loads (leading to a high probability of transmission per contact during the infectious period) but a shorter duration of infection. Given that we expect the trends for high contact densities to continue, we expect that the majority of viruses that emerge in human populations will continue to be those that cause acute infections. For animal diseases, a trend toward larger agricultural holdings that permit more contacts between animals may also lead to more acute viral infections in animals than observed currently.

In terms of evolutionary biology, maximizing “fitness” like *R*_0_ is used as a metric for evaluating the direction of evolution. For the sake of simplicity, in the current study, we considered only the evolution of the virus at an epidemiological equilibrium. We ignored the evolution of the host for simplicity, although we expect this to be reasonable because the generation time of a virus is much shorter than that of the host organism. However, to capture a long-term non-equilibrium evolution, it may be necessary to consider coevolutionary dynamics between hosts and viruses. A recent study by Sasaki et al. (2022) have shown that pathogens often showing antigenic escape from host immune defenses evolve through non-equilibrium dynamics, indicating that *R*_0_ is no longer the measure to understand viral evolutionary stable strategy [20]. Evolutionary game theory might be able to predict an evolutionarily stable strategy under the stochastic environmental scenario presented in this paper [45, 46].

In addition, our approach has several limitations. First, our mathematical model for individual-level virus infection, Eqs.(1–4), does not fully reflect the detailed physiologic processes of within-host virus infection. For example, we do not explicitly include a host immune response or antiviral treatment, both of which may affect the duration of the infectious period and peak viral load. However, we do not expect this to change our qualitative conclusions, as these effects may be mimicked by changing the values of the parameters of our model. Another modeling limitation is that our mathematical model for population-level transmission, Eqs.(7–8), does not account for “a transmission chain,” that is, we focused only on the sum of secondary cases generated throughout the infectious period by a primary case, *R*_0_. If entire chains of transmission are considered, it may be necessary to consider stochastic effects that may lead to particular viral phenotypes failing to become established in host populations, particularly in the region of a low-density contact history (the mean contact number from 0.01 to 1) for which *R*_0_ ≈ 1 [47]. Future studies may explore this further by nesting a viral dynamics model within a stochastic population-scale transmission model (using, for example, a stochastic adaptation of the multi-scale epidemiologic modeling framework proposed by [48]). Furthermore, explicitly considering transmission mode, host heterogeneity and detailed life history may also impact on the outcomes of viral evolution qualitatively.

Overall, we anticipate our study to be a key advance in modeling theory that associates the type of virus optimized to the host environment. We face emerging respiratory acute virus infections such as SARS-CoV-2, and host environments continue to change as transportation networks are developed and urban areas expanded. In particular, the frequency of contacts between people has increased yearly. As we showed here, such an environment favors acute viral infections. To prevent such emerging infectious diseases, it is crucial that infectious disease surveillance systems are strengthened, particularly in locations with high contact densities, allowing interventions to be introduced quickly to prevent the transmission of novel viruses and variants. This is of clear benefit for public health.

## Materials and Methods

To understand the dependence of viral evolution on the “environment,” we employed a multi-level population dynamics model (i.e., coupling a population-level virus transmission model and an individual-level virus infection model) and evaluated how the proliferative ability and accuracy of replication of viruses optimized to increase the reproductive number, *R*_0_ (see **Fig. 1A**). For the sake of simplicity, we considered an epidemiological equilibrium. We assumed that environments are characterized by the number of contacts per day (i.e., contact numbers among populations), and that an infected individual has a probability of infection (per contact with a susceptible individual) that depends on viral load. Note that the “noninfectious” period of infected individuals has not been directly accounted for in the infection probability (see below). We assumed that a newly infected individual has the potential to spread the infection to others immediately after infection, although the probability of infection is very small during this period. Below, we describe the details of the model as separate sections.

### Virus infection dynamics

We employed a simple mathematical model for virus infection dynamics considering the accuracy of viral replications [49]:

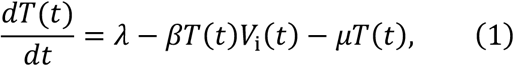

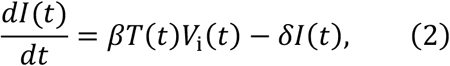

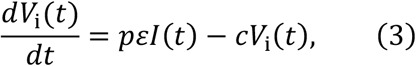

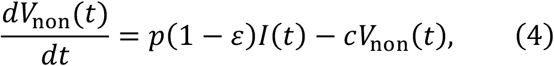

where the variables *T*(*t*), *I*(*t*), *V*_i_(*t*) and *V*_non_(*t*) are the number of uninfected target cells, infected target cells, and the amount of infectious virus (i.e., infectious viral load) and noninfectious virus at time *t* since infection, respectively. The parameters *β, δ, p*, and *c* represent the rate constant for virus infection, the death rate of infected cells, the per-cell viral production rate (i.e., proliferative ability of virus), and the per capita clearance rate of the virus, respectively. In addition, we assumed a fraction *ε* of the produced virus is infectious but 1 – *ε* is noninfectious (the value of *ε* represents the accuracy of replication). In other words, *pεI*(*t*) in this model represents ‘infectious virus production’, meaning the parameter *ε* includes influencing factors such as the accuracy of replication (which is the main factor and focused in this study), the accuracy of virion assembly, the rate of virus particle aggregation and others. To mimic viral cytopathogenesis depending on the viral replication level, we defined *δ* = *δ*_max_*p*/(*p* + *δ*_50_), that is, an increasing Hill function of *p*. The parameters *δ*_max_ and *δ*_50_ are the maximum value of *δ* and the viral production rate satisfying *δ* = *δ*_max_/2, respectively. We fixed the parameter values (except for *p* and *ε*) and initial values to correspond to biologically reasonable values or ranges (see below and **Table S1**). The sensitive analysis on *δ* is provided in **Fig.S5**.

### Duration of infectious period as a function of viral load

The duration of infectious period is assumed to depend on the total viral load (i.e., the cumulative viral load) as observed for several pathogens [50, 51]:

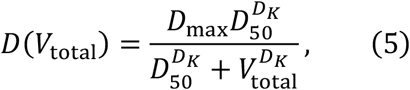

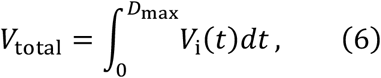

where *D*_max_ is the maximum duration of the infectious period, *D*_50_ is the viral load at which the duration is half of its maximum, and *D_k_* is the steepness at which duration decreases with increasing viral load [51].

### Probability of infection

To evaluate the daily transmissibility of an infected individual through the duration of infectious period, we assumed that for any given contact, each 10-fold increase in infectious viral load will lead to an r-fold increase in infectiousness [52]; thus the probability of infection *ρ* at time *t* is described by:

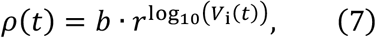

where *b* is the basilar probability of infection immediately after being infected with the virus.

### Number of encounters

To mimic a daily contact history, we assume the number of contacts on any day is drawn from a negative binomial distribution [53]. Since a negative binomial distribution is identical to a Poisson distribution where the mean parameter *λ* follows a gamma distribution, the daily contact numbers *C*(*t*) of a focal infectious individual at time *t* are generated by sampling from the following distributions (see **Fig.1C** as examples):

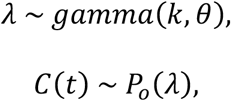

where *k* and *θ* are the shape parameter and the scale parameter, respectively. We note that *k* influences the skewness, and *θ* influences the variance of the distribution, respectively. The mean contact number through the history is the product of *k* and *θ*, that is, *kθ*.

As a supplementary analysis, we also considered a different approach for generating the contact numbers *C*(*t*), by considering a contact scenario on different networks: the Barabási–Albert (BA) scale-free network model and the Watts–Strogatz (WS) small-world network model (see **Text S2**, **Fig.S2**, **S3** and **Table S2**). The BA model generates a network such that a graph of *n* nodes is grown by attaching new nodes, each with *m* edges, that are preferentially attached to existing nodes with high degree [54]. The WS model generates a network such that starting from a ring over *n* nodes, which is connected with its *q* nearest neighbors, shortcuts are created by replacing some edges with probability *p* [55]. Contact numbers on a generated graph are calculated by counting the number of nodes that are connected with the one randomly chosen node on the network at each time step.

### Reproductive number and fitness landscape

We calculated the expected total number of secondary cases generated throughout the infectious period of a primary case (*R*_0_). On each day, the number of secondary cases is considered as the multiplication of the number of encounters (i.e., the contact number) and the probability of infection. Thus, *R*_0_ is calculated as below:

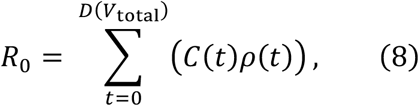

where *D*(*V*_total_) is the duration of infectious period of an infected individual defined by Eq.(5). Once fixing the daily contact number, *R*_0_ is dependent on the parameters appearing in the mathematical model for the virus infection dynamics, Eqs.(1–4). We hereafter focus on the dependence of the rate of proliferation and accuracy in the viral life cycle, *p* and *ε*, respectively. We calculate *R*_0_ in various patterns of (*p,ε*), and thus the fitness landscape of *R*_0_ is constructed as a function of *p* and *ε*.

### Evolution of virus to increase the reproductive number

One simple assumption is that viral population will eventually be dominated by the virus with the largest *R*_0_. From this perspective, a genetic algorithm [56], a population-based method of evolutionary computation, is implemented to search for the optimal point in the fitness landscape (i.e., maximizing *R*_0_) regarding the combination of *p* and *ε*. The genetic algorithm heuristically searches for the optimal *R*_0_ with a group of solutions made up of combinations of *p* and *ε* (see Alg.1 and Table S3 for details). Initially, solutions are randomly placed around the fitness landscape (Initialize) and are assigned *R*_0_ as fitness (Evaluate). Over successive iterations, the combinations of *p* and *ε* with higher *R*_0_ are selected to build the next solutions (selectSolutions) in a fitness proportionate manner where the solution with higher *R*_0_ has a greater chance to be chosen for the next iteration. Note that the two highest-performing solutions are always chosen for the next iteration (Elitism). To find better combinations of *p* and *ε*, solutions are recombined (crossover), and each parameter is slightly altered by adding a random variable drown from a Uniform distribution with a pre-determined range (Mutate). Finally, it searches a better solution in the expectation of finally converging to the combination of *p* and *ε* with the highest *R*_0_. These iterations can be considered as the virus “evolving” its properties of *p* and *ε* toward an optimal solution so that the virus can increase its own fitness in the evolutionary history.

#### Algorithm 1: Genetic Algorithm

~~~
Input: s_i_, solution i of combinations of *p* and *ε*
Input: P(t = i) = {s_1_, s_2_, ..., s_j_}, population of solutions in iteration i
t ← 0;
Initialize (P(t = 0)) with random *p* and *ε*;
Evaluate (P(t = 0));
while not termination do
      P(t)_e_ ← Elitism (P(t));
      P(t)_p_ ← selectSolutions (P(t));
      P(t)_c_ ← crossover (P(t)e +P(t)p);
      Mutate (P(t)c);
      Evaluate (P(t)c);
      P(t+1) . P(t)c;
      t ← t + 1;
end while
~~~

## Acknowledgments

This work was supported in part by a Grant-in-Aid for JSPS Scientific Research (KAKENHI) B 18H01139 (to S.I.), 16H04845 (to S.I.), Scientific Research in Innovative Areas 20H05042 (to S.I.); 21K15160 (to R.Y.), JP20H05682 (to K.T.), JP21K19909 (to K.T.); AMED CREST 19gm1310002 (to S.I.); AMED Development of Vaccines for the Novel Coronavirus Disease, 21nf0101638s0201 (to S.I.); AMED Japan Program for Infectious Diseases Research and Infrastructure, 20wm0325007h0001 (to S.I.), 20wm0325004s0201 (to S.I.), 20wm0325012s0301 (to S.I.), 20wm0325015s0301 (to S.I.); AMED Research Program on HIV/AIDS 22fk0410052s0401 (to S.I.); AMED Research Program on Emerging and Re-emerging Infectious Diseases 20fk0108140s0801 (to S.I.); AMED Program for Basic and Clinical Research on Hepatitis 21fk0210094 (to S.I.); AMED Program on the Innovative Development and the Application of New Drugs for Hepatitis B 22fk0310504h0501 (to S.I.); JST MIRAI JPMJMI22G1 (to S.I.); Moonshot R&D JPMJMS2021 (to S.I.) and JPMJMS2025 (to S.I.); Mitsui Life Social Welfare Foundation (to S.I.); Shin-Nihon of Advanced Medical Research (to S.I.); Suzuken Memorial Foundation (to S.I.); Life Science Foundation of Japan (to S.I.); SECOM Science and Technology Foundation (to S.I.); The Japan Prize Foundation (to S.I.); Foundation of Kinoshita Memorial Enterprise (to S.I.).

## Competing Interest Statement

The authors declare that they have no competing interests.

## Authors’ Contributions

Conceived and designed the study: RK AM KT WSH RNT SI RY. Analyzed the data: JS SI RY. Wrote the paper: All authors. All authors read and approved the final manuscript.

## Notes

### Competing Interest Statement

The authors have declared no competing interest.

