## supplementary material for "Contact-number-driven virus evolution: a multi-level modeling framework for the evolution of acute or persistent RNA virus infection"

^1^Department of Advanced Transdisciplinary Science, Hokkaido University, Sapporo, Hokkaido, Japan. ^2^Laboratory of RNA Viruses, Department of Virus Research, Institute for Life and Medical Sciences (LiME), Kyoto University, Kyoto, Japan. ^3^Mathematical Institute, University of Oxford, Oxford, United Kingdom. ^4^Mathematics Institute, University of Warwick, Coventry, United Kingdom. ^5^Zeeman Institute for Systems Biology and Infectious Disease Epidemiology Research, University of Warwick, Coventry, United Kingdom. ^6^Laboratory of RNA Viruses, Graduate School of Biostudies, Kyoto University, Kyoto, Japan. ^7^Department of Molecular Virology, Graduate School of Medicine, Kyoto University, Kyoto, Japan. ^8^interdisciplinary Biology Laboratory (iBLab), Division of Natural Science, Graduate School of Science, Nagoya University, Nagoya, Japan. ^9^Institute of Mathematics for Industry, Kyushu University, Fukuoka, Japan. ^10^Institute for the Advanced Study of Human Biology (ASHBi), Kyoto University, Kyoto, Japan. ^11^Interdisciplinary Theoretical and Mathematical Sciences Program (iTHEMS), RIKEN, Saitama, Japan. ^12^NEXT-Ganken Program, Japanese Foundation for Cancer Research (JFCR), Tokyo, Japan. ^13^Science Groove Inc., Fukuoka, Japan 8100041.

^1^To whom correspondence may be addressed.

 (S.I.) and (R.Y.).

Address: Furo-cho, Chikusa Nagoya 464-8602, Japan (S.I.), Phone: +81-52-789-2992 (S.I.) and Sapporo, Hokkaido 060-0810, Japan (R.Y.), Phone: +81-11-706-2659 (R.Y.)

**Supplementary text**

**Text S1. Difference in replication accuracy between persistently vs. acutely infectious viruses**

Replication accuracy will significantly impact transmission within a host population. In the case of HCV, experimental evidence suggests that a minor viral population with a much lower evolutionary rate than usual exists in persistently infected patients [1, 2]. The minor lineage avoid adaptive evolution within the host, is conserved as genome templates. Similarly, bovine viral diarrhea viruses (BVDV) strains that establish persistent infection evolve at a slower rate than acutely infected strains. Two biotypes of BVDV have been identified, the acutely infectious cp biotype and the persistently infectious ncp biotype [3]. The cp biotype is derived from the ncp biotype, and the cp biotype is a highly virulent strain that replicates quickly, has high viral load, and is lethal within weeks of infection. The genome sequences in persistently infected animals in the same herd have been found to be very similar [4-6]. In addition, sequence conservation of BVDV among offspring born from persistently infected animals is quite high, and no significant changes in BVDV genome sequences have been observed in persistently infected animals over a period of years [7]. As described above, epidemiological evidence has been presented that persistently infectious viruses have a lower evolutionary rate than acutely infectious viruses.

**Text S2. Virus evolution based on network-generating contact history**

We investigate how the properties of a virus evolve based on contact history generated from the following well-established network models: Barabási-Albert scale-free network model (BA model) and Watts-Strogatz small world network model (WS model). The contact number of a contact history is the number of nodes connected to a node chosen at random from all nodes in the network (see **Table S2** for the parameters used).

As shown by 50 independent iterations, a virus under the contact history generated by both the BA model and the WS model evolves high virus production but low accuracy (**Fig. S2C** and **Fig. S3C**). In both models, the median duration of the infectious period is 4.0 days (data not shown), which corresponds to the characteristics of the acute infection phenotype.

**Table S1**

**Parameters used for virus infection dynamics in the main text.**

| Parameter | value |
| --- | --- |
| Number of uninfected target cells at initial time, $\boldsymbol{T}\left( \boldsymbol{t=0} \right)$ | $1000$ |
| Number of infected target cells at initial time, $\boldsymbol{I}\left( \boldsymbol{t=0} \right)$ | $0$ |
| Amount of infectious virus at initial time, $\boldsymbol{V}_{\boldsymbol{i}}\left( \boldsymbol{t=0} \right)$ | $1$ |
| Amount of noninfectious virus at initial time $\boldsymbol{V}_{\mathbf{non}}\left( \boldsymbol{t=0} \right)$ | $0$ |
| Initial number of uninfected cells, $\boldsymbol{\lambda}$ | $10$ |
| Rate for virus infection, $\boldsymbol{\beta}$ | $0.0001$ |
| Death rate of noninfectious cells, $\boldsymbol{\mu}$ | $0.01$ |
| Death rate of infectious cells, $\boldsymbol{\delta}$ | $\delta_{\max}p/\left( p+\delta_{50} \right)$ |
| Maximum value of $\boldsymbol{\delta}$, $\boldsymbol{\delta}_{\mathbf{max}}$ | $1$ |
| Viral production rate satisfying $\boldsymbol{\delta}_{\mathbf{max}}\boldsymbol{/2}$, $\boldsymbol{\delta}_{\boldsymbol{50}}$ | $1000$ |
| Clearance rate of the virus, $\boldsymbol{c}$ | $10$ |
| Proliferative ability of virus, $\boldsymbol{p}$ | [0, 100, 200, …, 9900] (100 points in total) |
| Accuracy of virus replication, $\boldsymbol{\varepsilon}$ | [0.0, 0.01, 0.02, …, 0.99] (100 points in total) |
| Maximum duration of the infectious period, $\boldsymbol{D}_{\mathbf{max}}$ | $1000$ |
| Viral load at which the duration is half of its maximum, $\boldsymbol{D}_{\boldsymbol{50}}$ | $1.0\times{10}^{6}$ |
| Steepness at which duration decreases with increasing viral load, $\boldsymbol{D}_{\boldsymbol{k}}$ | $7.5$ |
| Basilar probability of infection immediately, $\boldsymbol{b}$ | $9.8\times{10}^{-8}$ |
| r-fold increase in infectiousness for each 10-fold increase in infectious viral load, $\boldsymbol{r}$ | $25$ |

**Table S2**

**Parameters used for the network models**

| model | parameter | value |
| --- | --- | --- |
| BA | number of nodes, $n$ | $100$ |
|  | number of edges to attach from a new node to existing nodes, $m$ | $1$ |
| WS | number of nodes, $n$ | $100$ |
|  | nearest nodes which are connected, $k$ | $2$ |
|  | the probability of rewiring each edge, $p$ | $0.5$ |

BA: Barabási-Albert scale-free network model

WS: Watts-Strogatz small world network model

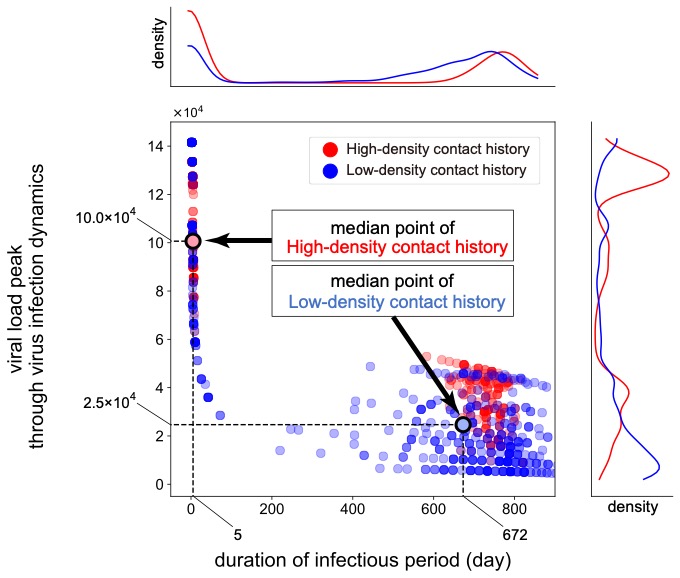

**Fig S1. Relation between duration of infectious period and peak viral load.** In each independent iteration, viruses evolve for the duration of the infectious period and the peak viral load depending on the contact history: high-density contact history (red) and low-density contact history (blue), respectively. The black arrows correspond to the median values representing the evolutionary endpoints. The parameter sets of shape and scale used are as follows: $\left( k, \theta\right)=\left( 1000, 0.002 \right)$ for high-density contact history, and $\left( k, \theta\right)=\left( 0.12, 10 \right)$ for low-density contact history, respectively. For both contact histories, the distributions of duration of infectious period are illustrated as kernel plots above the main figure. Also shown to the right of the main figure are the distributions of viral load peak through virus infection dynamics. For the high-density contact history, virus evolution results in a short duration of infectious period (5 days) with a high viral load ($1.0\times{10}^{5}$). In contrast, for low-density contact history, virus evolution results in a long duration of infectious period (672 days) with a low viral load ($2.5\times{10}^{4}$). 500 simulations were run for each contact history parameter set.

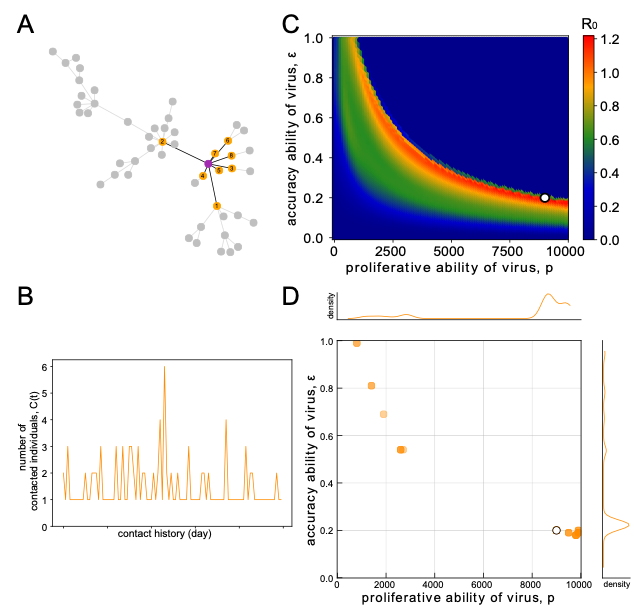

**Fig S2. Evolution of virus on a contact history generated from the** **Barabási-Albert scale-free network model.** An example graph of the BA model is illustrated in **(A)**. Contact numbers are calculated by counting the number of nodes that are connected to a single node chosen at random (purple). In this example, the contact number is 8 (orange). Iterating this process generates a contact history. An example of a given contact history is plotted in **(B)** (only 100 days is shown). Optimal parameter combinations ($p$ and $\varepsilon$) to increase $R_{0}$ are calculated for a given contact history generated from the BA model in **(C)**. Color represents the degree of $R_{0}$: blue corresponds to lower values of $R_{0}$, and red corresponds to higher values of $R_{0}$. The white dot represents the optimal point that increases $R_{0}$ in a single simulation (*see* the corresponding white dot in **(D)**). Using the same parameter set, 50 optima based on independent contact histories are calculated in **(D)**. The kernel plots illustrated at the top and right of the figure are the distribution of $p$ and $\varepsilon$, respectively.

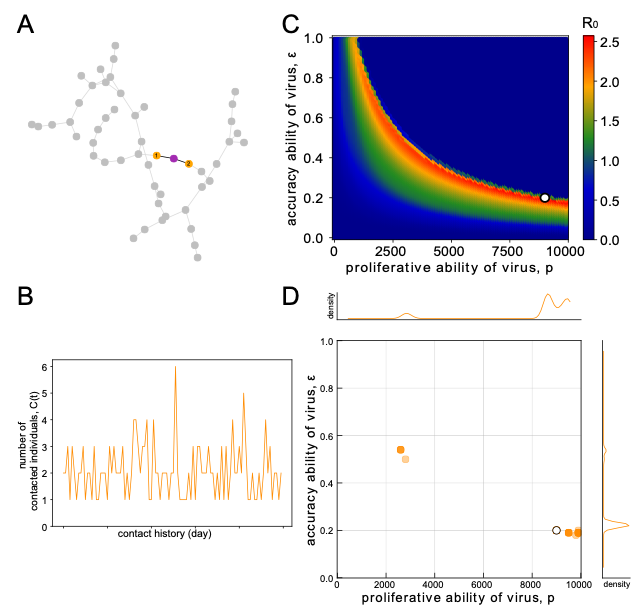

**Fig S3. Evolution of virus on a contact history generated from the Watts-Strogatz small world network model.** An example graph of the WS model is illustrated in **(A)**. Contact numbers are calculated by counting the number of nodes that are connected to a single node chosen at random (purple). In this example, the contact number is 2 (orange). Iterating this process generates a contact history. An example of a given contact history is plotted in **(B)** (only 100 days is shown). Optimal parameter combinations ($p$ and $\varepsilon$) to increase $R_{0}$ are calculated for a given contact history generated from the WS model in **(C)**. Color represents the degree of $R_{0}$: blue corresponds to lower values of $R_{0}$, and red corresponds to higher values of $R_{0}$. The white dot represents the optimal point that increases $R_{0}$ in a single simulation (*see* the corresponding white dot in **(D)**). Using the same parameter set, 50 optima based on independent contact histories are calculated in **(D)**. The kernel plot illustrated at the top and right of the figure are the distribution of $p$ and $\varepsilon$, respectively.

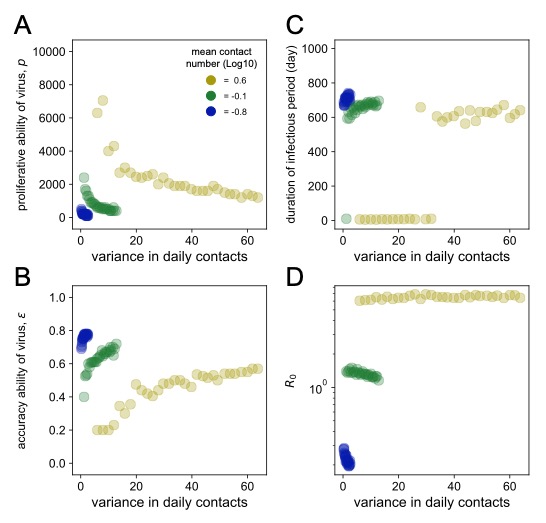

**Fig S4. Virus evolution in various scenarios of variance in daily contacts.** Each color represents the mean contact numbers of 0.6, -0.1, and -0.8, respectively, corresponding to the colored dots in **Fig.3** in the main text. We used the following parameter sets: the scale parameter $\theta$ ranges from 0.5, 1.0, 1.5, …, 14.5, 15.0 (30 points in total), and the shape parameter $k$ ranges to keep the specific mean contact number. We conducted a total of 500 independent iterations for each parameter set of $k$ and $\theta$, and showed the mean values. The combination of $p$ and $\varepsilon$ is plotted in **(A)** and **(B)**, separately. The duration of the infectious period and $R_{0}$ are plotted in (**C**) and (**D**), respectively.

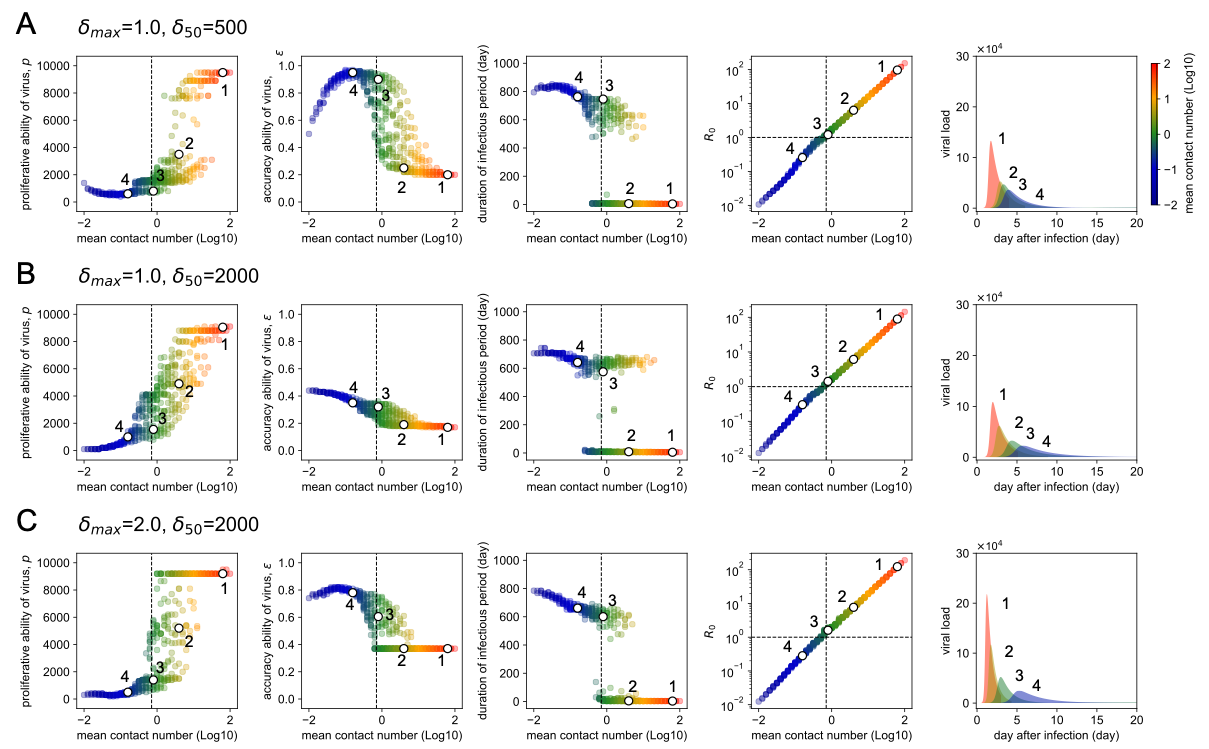

**Fig S5. Virus evolution in various scenarios of contact history.** Each row represents a single parameter set of ($\boldsymbol{\delta}_{\boldsymbol{max}}$ and $\boldsymbol{\delta}_{\boldsymbol{50}}$), that is **(A)**, **(B)**, **(C)** for (1.0, 500), (1.0, 2000), and (2.0, 2000), respectively. A total of 441 contact history patterns are explored for each parameter set of ($\boldsymbol{\delta}_{\boldsymbol{max}}$ and $\boldsymbol{\delta}_{\boldsymbol{50}}$), to determine the vial evolution to optimize $R_{0}$ in a given environmental scenario. We conducted a total of 500 independent iterations for each parameter combination of $k$ and $\theta$ as well as in **Fig.3** in the main text. Numbers 1, 2, 3, and 4 in the panels of viral load are the corresponding dynamics illustrated as white dots in the columns of $p$, $\varepsilon$, the duration of infectious period, and $R_{0}$, respectively. In the panels of $R_{0}$, the vertical dashed line separates the area of mean contact number by an index, $R_{0}={10}^{0} \left( =1 \right)$ described by the horizontal dashed lines. The corresponding dashed line is also illustrated in the panels of $p$, $\varepsilon$, and the duration of infectious period.
